# Tetrameric TRF2 forms t-loop to protect telomeres from ATM signaling and cNHEJ

**DOI:** 10.1101/2024.06.27.600884

**Authors:** Audrey M. Goldfarb, Nanda K. Sasi, Teague C. Dilgen, Sarah W. Cai, Logan R. Myler, Titia de Lange

## Abstract

Telomeres often occur in the t-loop structure, formed through the base-pairing of the 3’ telomeric overhang is base-paired with more internal telomeric DNA. T-loops are proposed to avert ATM kinase signaling and classical non-homologous end joining (cNHEJ) by sequestering the telomere end. The shelterin subunit TRF2 is required for t-loop formation but how it generates t-loops is not known and the t-loop model for telomere protection remains untested. We show that TRF2 binds telomeric DNA as a tetramer using two dimerization domains and that tetramerization is critical for telomere protection. Furthermore, TRF1, the TRF2 paralog that binds DNA as a dimer and does not form t-loops, gained the ability to form t-loops in vivo upon its tetramerization through the addition of a second dimerization domain. We propose that TRF2 tetramers loop telomeric DNA, resulting in torsional stress and opening of the helix, which is a requirement for t-loop formation. Importantly, t-loops formed by tetrameric TRF1 were sufficient to avert ATM signaling and blocked most cNHEJ at telomeres, demonstrating that t-loops per se protect telomeres from these threats.

---

How telomeres solve the end-protection problem is a fundamental question with implications for human health as loss of telomere protection is central to telomere biology disorders (*1*) and the telomere tumor suppressor pathway (*2*). Human telomeres are bound by shelterin, a six-subunit protein complex that is critical for telomere protection (*3*). Vertebrate shelterin complexes contain two related proteins that bind to the double-stranded (ds) TTAGGG repeat array, TRF1 and TRF2 (*4–6*). Both have C-terminal Myb/SANT DNA binding domains and a large dimerization domain, the TRF homology (TRFH) domain, but their functions have diverged. Whereas deletion of TRF2 results in activation of ATM kinase signaling at telomeres and their fusion by cNHEJ (*7–11*), deletion of TRF1 does not affect telomere protection but results in defects in the replication of telomeric DNA (*12–16*). Furthermore, TRF2, but not TRF1, is necessary and sufficient to form and/or stabilize t-loops (*17*, *18*), the large lariats that are formed when the 3’ overhang of the telomere base-pairs with a more internal part of the telomeric repeat array (*19*). Based on these observations it was proposed that t-loop formation by TRF2 is the key mechanism by which telomeres are protected from ATM signaling and cNHEJ.

Although several findings are consistent with this proposal (e.g., (*20*, *21*), concerns about the model have been raised. First, TRF2 can repress cNHEJ using mechanisms that do not involve t-loop formation (*21–26*). Second, t-loops have been detected at less than 50% of telomeres, although this may be due to technical limitations (*17–19*, *21*). Third, in mouse embryonic stem (ES) cells, t-loops are formed in the absence of TRF2 (*27*, *28*). As noted previously (*3*), a strict test of the t-loop model for telomere protection would require an orthogonal, TRF2-independent method for generating telomeres with t-loops.Here we reveal the mechanism by which TRF2 forms t-loops, engineer a version of TRF1 that can generate t-loops in vivo, and use this use this form of TRF1 to show that t-loops protect telomeres from both ATM signaling and cNHEJ.

## Results

### DNA wrapping mutations do not interfere with telomeric protection by TRF2

Elegant biochemical studies suggested a compelling model for how TRF2 generates t-loops (*29*). It was shown that TRF2 can wrap the telomeric DNA around its TRFH dimerization domain, inducing topological stress that was proposed to lead to helix unwinding, thereby promoting the strand invasion that establishes the t-loop configuration (Fig. 1A). Alanine substitutions of 7 lysine and 2 arginine residues involved in DNA wrapping resulted in a version of TRF2, called TRF2-Topless, that lacked DNA wrapping activity in vitro and showed reduced t-loop formation and diminished protection from ATM signaling in vivo. However, additional data suggested that mouse TRF2-Topless localized to telomeres less efficiently than wild type TRF2 (*18*), confounding the interpretation of its telomere protection defect.

**Fig. 1.**
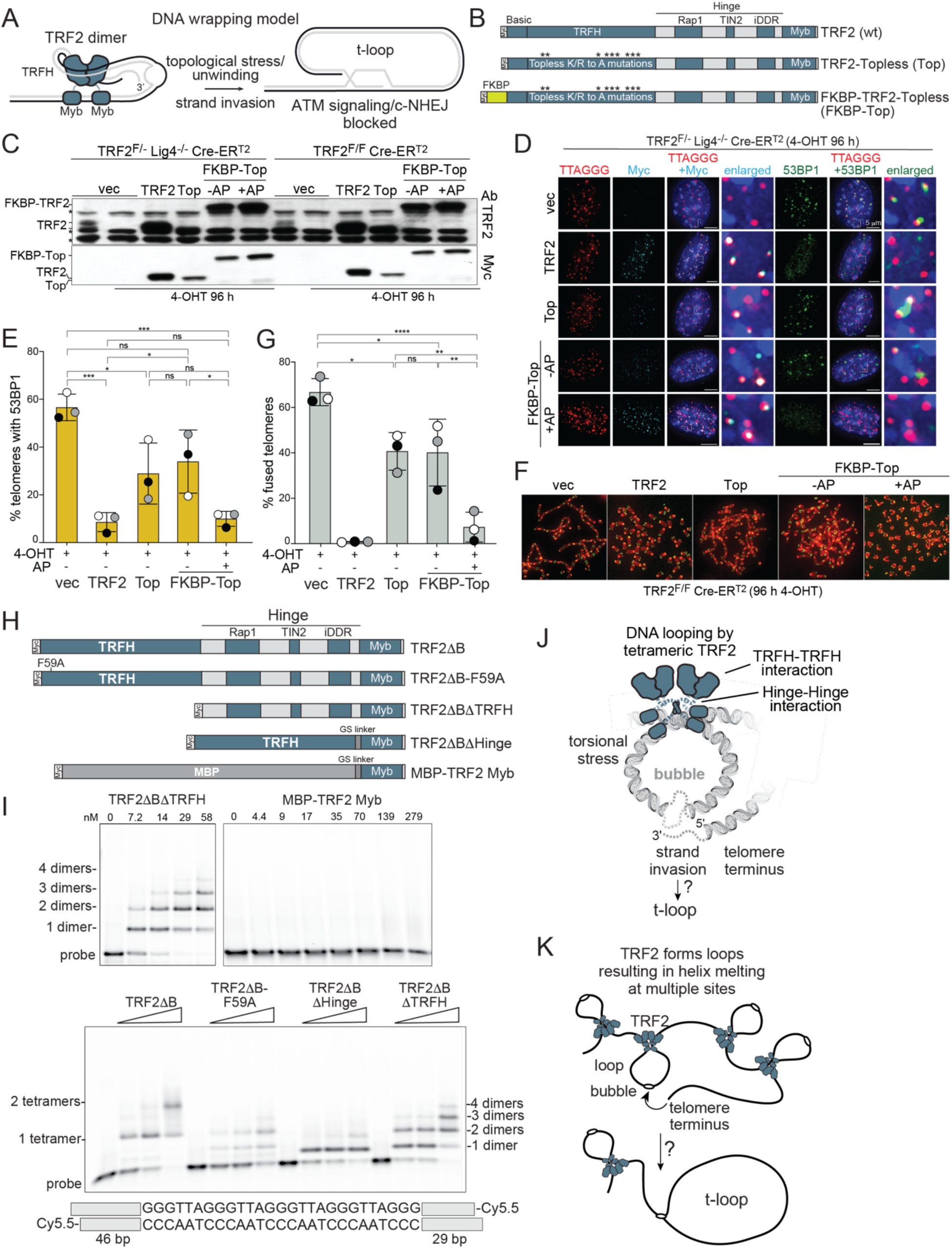
Testing DNA binding features of TRF2 relevant to t-loop formation. (**A**) Proposed model for t-loop formation based on the DNA wrapping activity of the TRF2 TRFH domain. (**B**) Schematic of Myc-tagged wild type (wt) TRF2, TRF2-Topless (Top), and FKBP-TRF2-Topless (FKBP-Top). Asterisks: Topless K-to-A and R-to-A mutations in the TRFH domain that interfere with DNA wrapping. (**C**) Immunoblots for the indicated TRF2 proteins expressed in TRF2^F/-^ Lig4^-/-^ Cre-ER^T2^ MEFs and TRF2^F/F^ Cre-ER^T2^ MEFs detected with an antibody to mouse TRF2 and Myc. Asterisks: nonspecific bands. Cre was induced with 4-OHT for 8 h and cells were harvested 96 h after 4-OHT addition. +AP: addition of AP1903 at 72 h before harvest. (**D**) Examples of the localization of the indicated Myc-tagged TRF2 proteins (IF, cyan) and 53BP1 (IF, green) at telomeres detected with TTAGGG repeat FISH (red). DNA stained with DAPI. Merged and enlarged images are shown. Induction of Cre and AP-mediated FKBP dimerization as in (C). (**E**) Quantification of TIF response as detected in (D). Data represent means ±SDs of three biological replicates of 30 nuclei each. ns, P>0.05; *, P<0.05; **, P<0.01; ***, P<0.001; ****, P<0.0001. P values based on ordinary one-way ANOVA with multiple comparisons. (**F**) Detection of telomere fusions in metaphase spreads from the indicated cells. DNA stained with DAPI (red), and telomeres detected with FISH (green). (**G**) Quantification of the percentage of telomeres that are fused detected as in (F). Data represent means ±SDs of three biological replicates with 20 metaphases each. ns, P>0.05; *, P<0.05; **, P<0.01; ***, P<0.001; ****, P<0.0001. P values based on ordinary one-way ANOVA with multiple comparisons. (**H**) Schematic of TRF2ΔB (lacking the Basic domain), TRF2ΔB-F59A, TRF2ΔBΔTRFH, TRF2ΔBΔHinge, TRF2ΔBΔHinge-F59A, and MBP-TRF2 Myb. The F59A mutation in helix 1 of the TRFH is predicted to disrupt dimerization. In MBP-TRF2 Myb, the 42.5 kDa monomeric Maltose Binding Protein (MBP) is fused to the Myb domain of TRF2. MBP-TRF2 Myb is 48 kDa and TRF2ΔB is 51 kDa. (**I**) EMSAs with the indicated TRF2 proteins schematized in (H) purified from E. coli. A schematic of the Cy5.5-labeled telomeric DNA probe is given below the gel. Upper gels: protein concentrations indicated. Lower gel: protein concentrations were: 30, 60, 120 nM. The probe concentration was 0.5 nM. Note that the smallest complex formed by TRF2ΔB is consistent with a tetramer (two dimers) whereas the smallest complex formed by TRF2ΔB-F59A represents a dimer. Monomeric TRF proteins do not form a complex in EMSAs. **(J)** Schematic highlighting the inferred dimerization interfaces that allow TRF2 to form a tetramer. (**K**) Model for t-loop formation by tetrameric TRF2. Binding of a TRF2 tetramer to DNA causes looping and torsional stress. The resulting unwinding of the duplex DNA promotes strand invasion of the 3’ overhang.

We found that the telomeric localization of TRF2-Topless was improved by tagging the N-terminus with the F36V variant of FKBP (*30*) and induction of FKBP dimerization in the presence of AP1903 (Fig. 1B-D). Consistent with previous reports, in TRF2^F/-^ Lig4^-/-^ mouse embryo fibroblasts (MEFs) from which TRF2 was deleted with Cre, neither TRF2-Topless nor the FKBP-tagged version fully repressed ATM kinase signaling as inferred from the accumulation of 53BP1 at telomeres (referred to as telomere dysfunction induced foci, TIFs (*31*)). As expected, ectopic wild type TRF2 reduced the TIF response to background levels (Fig. 1D, E). Importantly, the addition of AP1903 allowed FKBP-TRF2-Topless to repress the DNA damage signal to nearly the same extent as wild type TRF2 (Fig. 1D, E). In the cNHEJ-deficient Lig4^-/-^ MEFs chosen for in these experiments, the telomeres do not fuse, thereby avoiding the confounding effects of telomere fusions extinguishing the TIF response. In Cre-treated c-NHEJ-proficient TRF2^F/F^ MEFs, FKBP-TRF2-Topless was able to protect telomeres from end-to-end fusions in the presence of AP1903 but not in its absence (Fig. 1F, G). These data suggest that the topological stress induced by the DNA wrapping around the TRFH domain of TRF2 is not the only mechanism by which TRF2 protects telomeres.

### TRF2 binds DNA as a tetramer

TRF1 was shown to bind telomeric DNA as a dimer (*32–34*) and it was assumed that TRF2 similarly binds DNA as a dimer. However, atomic force microscopy (AFM) experiments suggested that TRF2, but not TRF1, can form a larger complex on telomeric DNA, potentially representing a tetramer (*34*). We determined how TRF2 binds telomeric DNA using the electrophoretic mobility shift assay (EMSA) with recombinant proteins and a telomeric repeat probe that can bind four Myb domains (Fig. 1H, I; Fig. S1A-E). The tested versions of TRF2 lacked the N-terminal Basic domain (Fig. 1H, Fig. S1A, B), which is not required for t-loop formation in vivo (*35*) and can lead to trapping of the DNA-protein complex in the slots of EMSA gels (*5*).

Unexpectedly, a version of TRF2 lacking its TRFH dimerization domain bound the telomeric DNA probe (Fig. 1H, I; Fig. S1C). Since a monomeric form of the TRF2 Myb domain (MBP-fused Myb) did not yield a detectable DNA-bound complex, the DNA binding by TRF2ΔTRFH suggests that it is a dimer. If TRF2ΔTRFH can form a dimer, full length TRF2 is expected to form a tetramer. Consistent with this prediction, TRF2-F59A which has the same size as wild type TRF2 but has a point mutation that diminishes the TRFH-TRFH interaction (Fig. S1D), formed a complex with DNA that was smaller than wild type TRF2 (Fig. 1I; Fig. S1C). TRF2-F59A also formed a second complex, likely representing two dimers bound to the probe. This second TRF2-F59A complex co-migrated with the wild type TRF2 complex (Fig. 1I; Fig. S1C), consistent with wild type TRF2 forming a tetramer. Formation of TRF2 tetramers and TRF2ΔTRFH dimers indicated that the second dimerization interface of TRF2 involves the Hinge region. Indeed, TRF2 lacking Hinge region formed DNA-bound complexes consistent with a dimeric protein (Fig. 1I; Fig. S1C), and the DNA binding of this deletion mutant was diminished by the F59A mutation in the TRFH (Fig. S1C-E).

The EMSA results were confirmed with TRF2 mutants produced by in vitro transcription/translation in a rabbit reticulocyte lysate (Fig. S2A-C). TRF2ΔHinge and TRF2ΔTRFH bound DNA as a dimers whereas TRF2 bound primarily as a tetramer. As was the case with MBP fused to the TRF2 Myb domain, a fusion of the TRF2 Myb domain to the TRFH of TIN2, which does not form dimers (*36*), resulted in a protein with no detectable DNA binding activity (Fig. S2A-C). Consistent with TRF2 but not TRF1 having a second dimerization domain, TRF2ΔTRFH had the ability to accumulate at telomeres whereas TRF1ΔTRFH did not (Fig. S3A-C).

The finding that TRF2 binds DNA as a tetramer – whereas TRF1 does not – suggests a mechanism that allows TRF2, but not TRF1, to form t-loops. A tetrameric TRF2 could loop telomeric DNA as has been observed by EM and AFM analysis (*34*, *37*). Like DNA wrapping, DNA looping can induce torsional stress and unwinding of the DNA helix, promoting the invasion of the 3’ overhang (Fig. 1J, K).

### TRF2 requires two dimerization domains for telomere protection

If tetramerization of TRF2 is critical for telomere protection, it is predicted that deletion of its Hinge domain, which has no effect on the telomeric localization of TRF2 (Fig. S3), would diminish telomere protection and that the function of such deletion mutants can be restored by orthogonal dimerization, e.g., FKBP in the presence of AP1903. Because deletion of the Hinge domain could potentially generate a dominant-negative version of TRF2, TRF2ΔHinge was expressed under the control of the Tet promoter and induced after deletion of the endogenous TRF2 (Fig. 2A, B). Upon induction of their expression, both the TRF2ΔHinge mutant and the FKBP-tagged version of this mutant were deficient in repression of ATM signaling at telomeres (Fig. 2C). Importantly, the protective function of FKBP-TRF2ΔHinge was largely restored by addition of AP1903 (Fig. 2C, D). Similarly, when these versions of TRF2 were induced in cNHEJ-proficient cells, the mutants lacking the Hinge showed a defect in cNHEJ inhibition and the block to cNHEJ could be partially restored by addition of AP1903 to cells expressing FKBP-TRF2ΔHinge (Fig. 2E-G). A confounding issue with the TRF2ΔHinge mutant is that this version of TRF2 gives rise to telomere connections that are detectable by both telomeric FISH and IF for TRF2 (Fig. 2C). As the molecular nature of these structures is unknown, we cannot exclude that the unusual structure of the telomeres affect ATM kinase signaling and cNHEJ. Nonetheless, the data are consistent with the notion that TRF2 is more effective at protecting telomeres when it can form a tetramer.

**Fig. 2.**
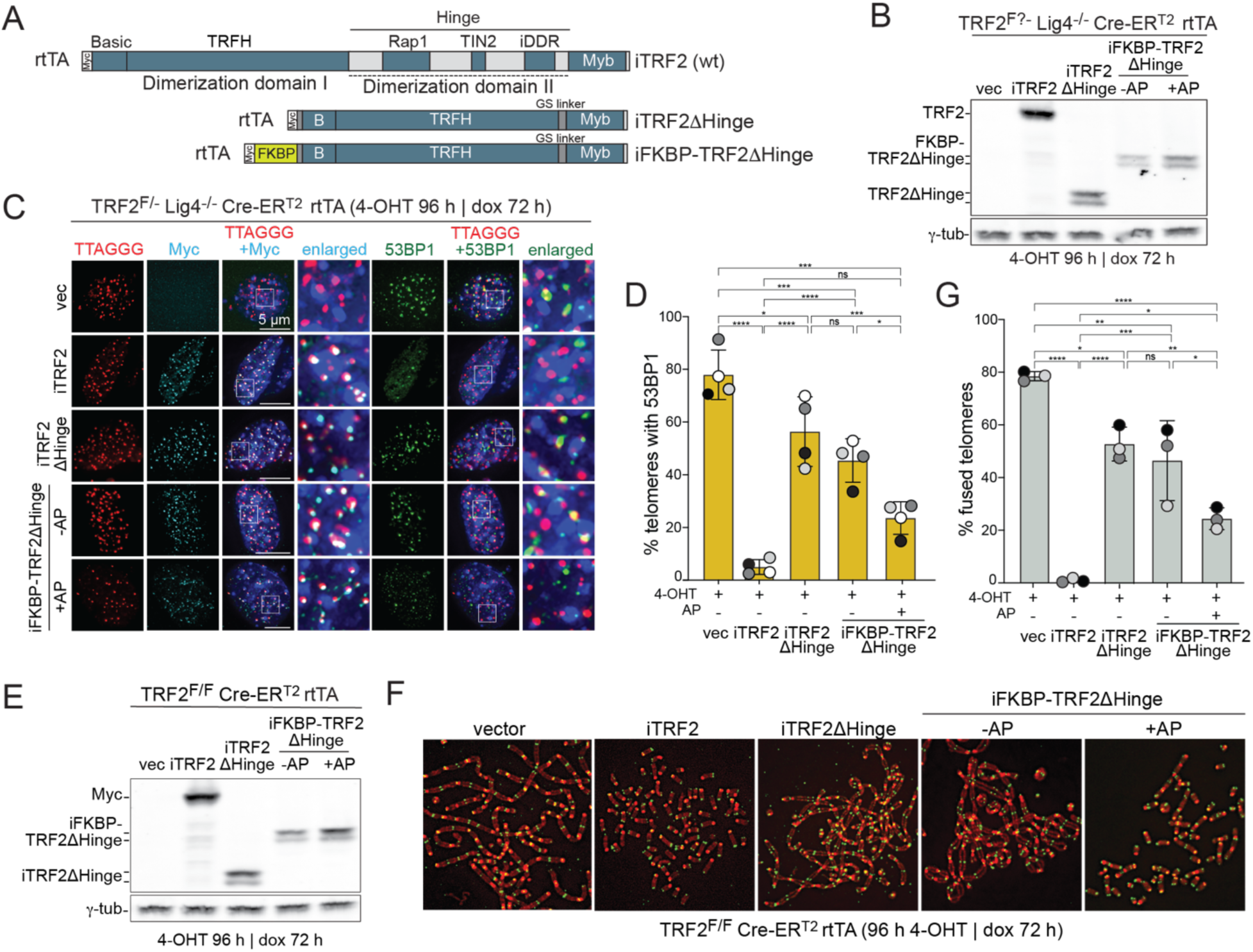
TRF2 requires two dimerization domains for telomere protection. (**A**) Schematic of Myc-tagged wt TRF2, TRF2τιHinge, and FKBP-TRF2τιHinge. Relatively equal levels of proteins were expressed from a Dox-inducible promoter using different doxycycline concentrations (TRF2: no Dox; TRF2τιHinge and FKBP-TRF2τιHinge:1 μg/mL). (**B**) Immunoblots for the indicated TRF2 proteins expressed in TRF2^F/-^ Lig4^-/-^ Cre-ER^T2^ MEFs detected with an antibody to Myc. Asterisks: nonspecific bands. Cre was induced with 4-OHT for 8 h and cells were harvested 96 h after 4-OHT addition. Expression of TRF2 proteins was induced with Dox for 72 h before harvest. +AP: addition of AP1903 at 72 h before harvest. (**C**) Examples of the localization of the indicated Myc-tagged TRF2 proteins (IF, cyan) and 53BP1 (IF, green) at telomeres detected with TTAGGG repeat FISH (red). DNA stained with DAPI. Merged and enlarged images are shown. Induction of Cre, Doxycycline, and AP-mediated FKBP dimerization as in (B). (**D**) Quantification of TIF response as detected in (C). Data represent means ±SDs of three biological replicates of 30 nuclei each. ns, P>0.05; *, P<0.05; **, P<0.01; ***, P<0.001; ****, P<0.0001. P values based on ordinary one-way ANOVA with multiple comparisons. (**E**) Immunoblots for the indicated TRF2 proteins expressed in TRF2^F/F^ Cre-ER^T2^ MEFs detected with an antibody to Myc. Asterisks: nonspecific bands. Cre was induced with 4-OHT for 96 h. Expression of TRF2 proteins was induced with Dox for before 72 h. +AP: addition of AP1903 at 72 h before harvest. (**F**) Detection of telomere fusions in metaphase spreads from the indicated cells. DNA stained with DAPI (red) and telomeres detected with FISH (green). (**G**) Quantification of the percentage of telomeres that are fused detected as in (F). Data represent means ±SDs of three biological replicates of 20 metaphase spreads each. ns, P>0.05; *, P<0.05; **, P<0.01; ***, P<0.001; ****, P<0.0001. P values based on ordinary one-way ANOVA with multiple comparisons.

### Tetrameric TRF1 can form t-loops and protect telomeres

To test the tetramerization model more stringently, we aimed to convert the dimeric TRF1 into a tetrameric protein by tagging its N-terminus with FKBP and adding AP1903 (Fig. 3A-C). TRF1, FKBP-TRF1, and FKBP-TRF1ΔTRFH were expressed in TRF2^F/-^ Lig4^-/-^ MEFs alongside TRF2 as a positive control. After deletion of the endogenous TRF2, the frequency of telomeres in the t-loop configuration was determined by Structured Illumination Microscopy (SIM) imaging of psoralen-crosslinked DNA spreads (*17*) (Fig. 3D, E). As expected from prior work (*17*, *18*, *21*, *27*, *35*), the frequency of telomeres in the t-loop configuration was reduced when TRF2 was absent and expression of exogenous TRF2 rescued this phenotype. TRF1 and FKBP-tagged TRF1 did not generate t-loops in cells lacking TRF2, consistent with TRF2 being the only subunit in shelterin mediating t-loop formation (*17*, *18*). However, when FKBP-tagged TRF1 was induced to form tetramers by the addition AP1903, the frequency of t-loops was indistinguishable from that in TRF2-proficient cells (Fig. 3D, E). Importantly, this t-loop forming activity of FKBP-TRF1 was dependent on its TRFH domain, consistent with the idea that FKBP-TRF1 can only form t-loops when it binds to telomeres as a tetramer. In the absence of AP1903, FKBP-TRF1 appeared to promote t-loop formation slightly better than TRF1, although this difference was not statistically significant (Fig. 3E). It is possible that FKBP-tag affects the behavior of TRF1, perhaps generating unstable tetramers. Consistent with this possibility, unlike TRF1ΔTRFH (Fig. S3A-C), FKBP-TRF1ΔTRFH showed some telomere localization even in the absence of AP1903 (see arrowheads in Fig. 3F), although its telomeric localization of was strongly improved by the dimerizer.

**Fig. 3.**
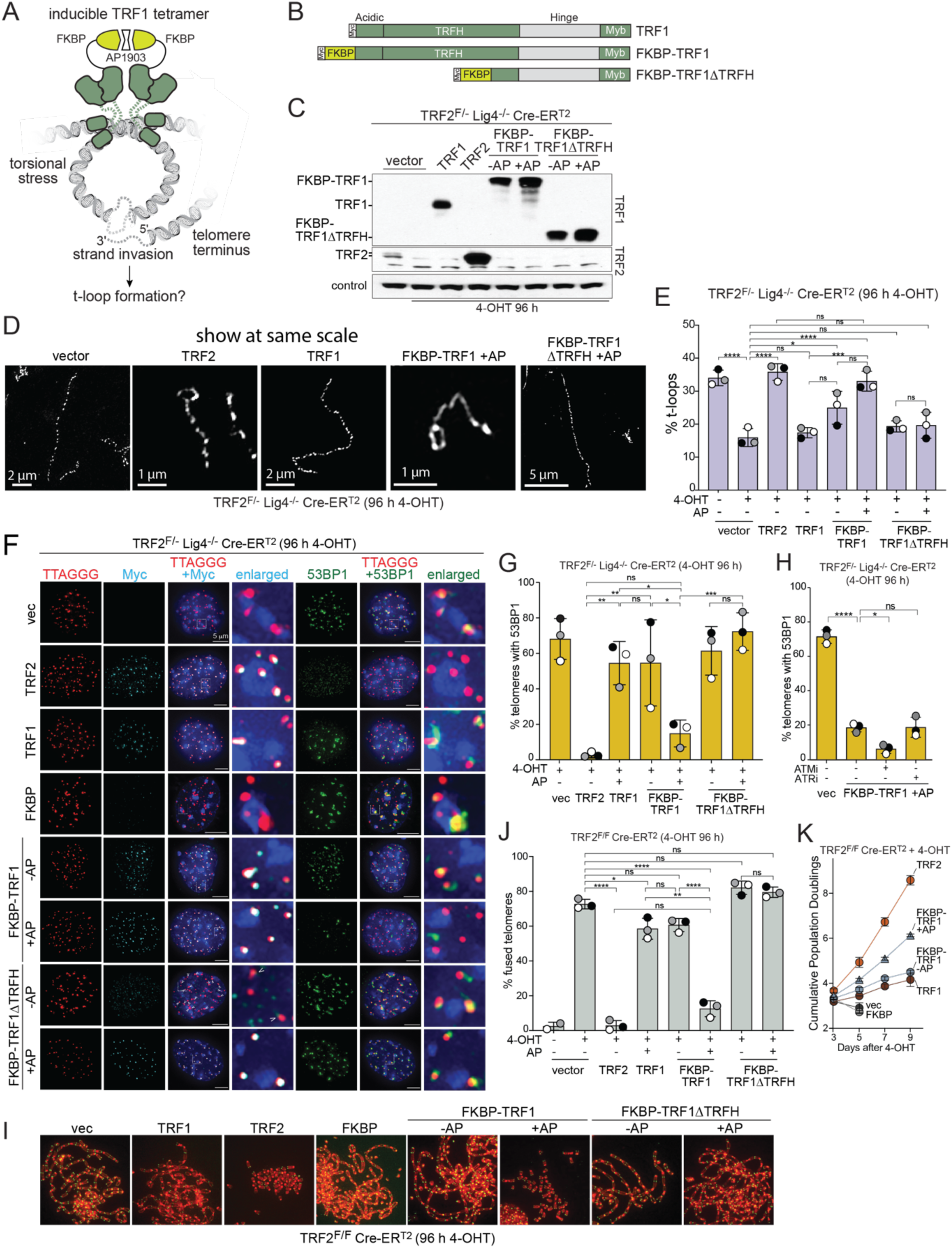
Tetrameric TRF1 forms t-loops and protects telomeres. (**A**) Model of t-loop formation by tetrameric TRF1. (**B**) Schematics of Myc-tagged TRF1, FKBP-TRF1, and FKBP-TRF1ΔTRFH. (**C**) Immunoblots for the indicated proteins expressed in TRF2^F/-^ Lig4^-/-^ Cre-ER^T2^ MEFs and TRF2^F/-^ Cre-ER^T2^ MEFs detected with an antibody to mouse TRF2 and TRF1. Cre was induced with 4-OHT for 8 h and cells were harvested 96 h after 4-OHT addition. +AP: addition of AP1903 at 72 h before to harvest. (**D**) Detection of t-loop formation by super-resolution SIM imaging on psoralen-crosslinked chromatin from indicated cells. Telomeric DNA was identified by FISH. (**E**) Quantification of t-loop frequency determined as in (D). Data represent means ±SDs of three biological replicates. ns, P>0.05; *, P<0.05; **, P<0.01; ***, P<0.001; ****, P<0.0001. P values based on ordinary one-way ANOVA with multiple comparisons. (**F**) Examples of the localization of the indicated Myc-tagged proteins (IF, cyan) and 53BP1 (IF, green) at telomeres detected with TTAGGG repeat FISH (red). DNA stained with DAPI. Merged and enlarged images are shown. Induction of Cre and AP-mediated FKBP dimerization as in (B). (**G**) Quantification of TIF response as detected in (F). Data represent means ±SDs of three biological replicates of 30 nuclei each. ns, P>0.05; *, P<0.05; **, P<0.01; ***, P<0.001; ****, P<0.0001. P values based on ordinary one-way ANOVA with multiple comparisons. (**H**) Quantification of TIF response in TRF2^F/-^ Cre-ERT Lig4^-/-^ MEFs infected with FKBP-TRF1 and treated with 4-OHT (96 h), AP1903 (72 h), and either ATMi (5 μM, 24 h) or ATRi (1 μM, 24 h). Data represent means ±SDs of three biological replicates of 30 nuclei each. ns, P>0.05; *, P<0.05; **, P<0.01; ***, P<0.001; ****, P<0.0001. P values based on ordinary one-way ANOVA with multiple comparisons. (**I**) Detection of telomere fusions in metaphase spreads from the indicated cells. DNA stained with DAPI (red) and telomeres detected with FISH (green). (**J**) Quantification of the percentage of telomeres that are fused detected as in (I). Data represent means ±SDs of three biological replicates of 20 metaphase spreads each. ns, P>0.05; *, P<0.05; **, P<0.01; ***, P<0.001; ****, P<0.0001. P values based on ordinary one-way ANOVA with multiple comparisons. (**K**) Growth curves of TRF2^F/F^ Cre-ERT MEFs treated with 4-OHT (8 h) at Day 0. +AP condition was treated with AP1903 throughout.

In agreement with the data on t-loop formation, FKBP-TRF1 largely restored the repression of ATM signaling at telomeres of Cre-treated TRF2^F/-^ Lig4^-/-^ cells when AP1903 was supplied. The same result was obtained with TRF2^F/F^ Ku70^-/-^ cells (Fig. S4A-C). The residual level of 53BP1 TIF formation in the presence of tetrameric TRF1 was due to ATM signaling as shown by treatment with ATM and ATR inhibitors (Fig. 3H). This low level of ATM signaling is expected from the inability of TRF1 to recruit the Apollo nuclease which only binds to the TRFH domain of TRF2 (*38*). In absence of Apollo, telomeres that have been replicated by leading-end DNA synthesis remain blunt and activate the ATM kinase (*39–43*), unless alternative resection pathways generate the 3’ overhang needed for t-loop formation. As was the case for t-loop formation, the ability of FKBP-TRF1 to protect telomeres from ATM signaling in the presence of AP1903 was dependent on the TRFH domain of TRF1 (Fig. 3G). Similar to the repression of ATM signaling, FKBP-TRF1 repressed cNHEJ at telomeres in the presence of AP1903 and this activity was again dependent on the TRFH domain of TRF1 (Fig. 3I, J). Controls with a TRF1 fused to a version of FKBP that does not bind AP1903 (*30*) further confirmed that formation of a TRF1 tetramer was responsible for telomere protection (Fig. S4D-H). Finally, the ability of FKBP-TRF1 to protect telomeres in the presence of AP1903 led to a substantial improvement of the proliferation of cells lacking TRF2 (Fig. 3K). These data, together with the data on FKBP-TRF2ΔHinge are consistent with the proposal that a tetrameric form of either TRF1 or TRF2 can form t-loops and thereby protect telomeres.

### t-loops repress cNHEJ

The fusion of dysfunctional telomeres by cNHEJ is dependent on their increased mobility, which is mediated by 53BP1 (*44*, *45*). As telomeres with tetrameric TRF1 do not activate ATM signaling and hence lack 53BP1, the absence of telomere fusions in this setting does not inform on the ability of t-loops to block cNHEJ. In prior work, Cre-treated TRF2^F/-^ ATM^-/-^ cells could be induced to join their telomeres by enforcing 53BP1 accumulation through local activation of ATR signaling (*46*). We therefore tested whether depletion of TPP1, which is needed for repression of ATR signaling by POT1a and POT1b (*47*), leads to fusion of telomeres that are protected by tetrameric TRF1. As expected, an shRNA to TPP1 induced a TIF response in cells containing either TRF2 or the tetrameric form of TRF1 (Fig. 4A-C). The TPP1 shRNA also induced a low level of telomere fusions in TRF2^F/F^ MEFs not treated with Cre (Fig. 4D, E; white arrows), consistent with prior findings (*36*, *46*, *47*). When TPP1 was depleted in TRF2-deficient cells expressing tetrameric TRF1, telomere fusions were observed (Fig. 4D, E). However, these fusions appeared to be largely due to alt-EJ rather than cNHEJ since they were dependent on PARP activity (Fig. 4D, E). The activation of alt-EJ at telomeres deprived of TPP1 is consistent with prior work indicating that alt-EJ is repressed by POT1a and POT1b (*36*, *48*) (Fig. 4D, E). In contrast, PARP inhibition did not reduce the telomere fusions induced by TRF2 deletion, which is a setting where the end-joining is mediated by cNHEJ (Fig. 4D, E). We further tested whether the shTPP1-induced telomere fusions were independent of cNHEJ using TRF2^F/-^ Lig4^-/-^ cells expressing induced tetrameric FKBP-TRF1. As expected, knockdown of TPP1 also gave rise to a telomere damage signal and telomere fusions, further confirming that the joining of telomeres is primarily due to alt-EJ (Fig. 4F-H; Fig. S5A, B). Since most, if not all, joining of telomeres bearing tetramerized FKBP-TRF1 were mediated by alt-EJ, we conclude that the t-loops formed by tetrameric TRF1 repress both ATM signaling and cNHEJ.

**Fig. 4.**
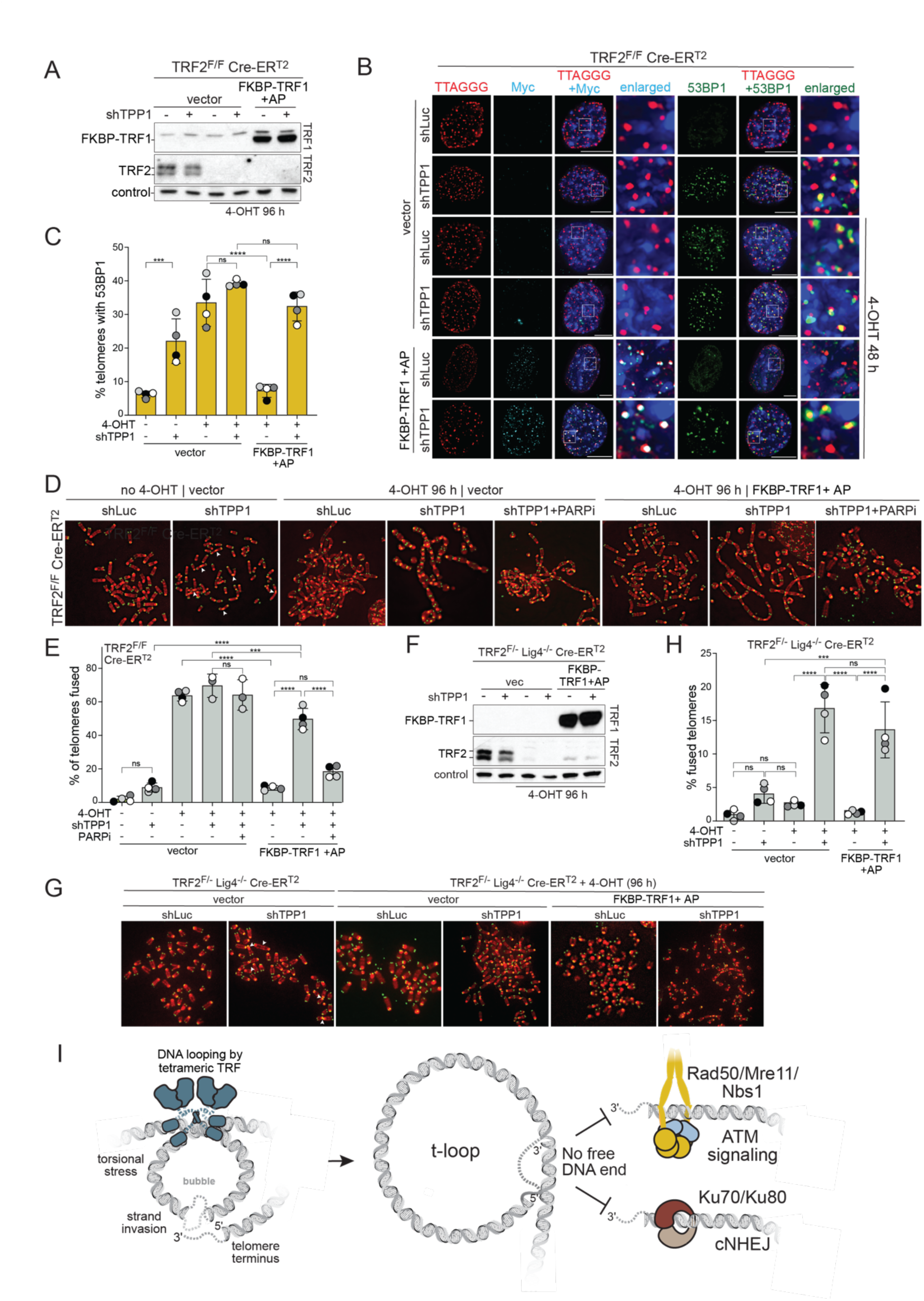
T-loops suppress cNHEJ-mediated telomere fusions. (**A**) Immunoblots for FKBP-TRF1 expressed in TRF2^F/F^ Cre-ER^T2^ MEFs detected with an antibody to mouse TRF1. Cre was induced with 4-OHT, and TRF2 knockout was confirmed 96 h after 4-OHT addition with an antibody against mouse TRF2. +AP: addition of AP1903 at 72 h before harvest. +shTPP1: cells were infected thrice with a shTPP1 or shLuc retroviral vector. The second transduction was carried out at t=0 and coincided with 4-OHT addition. The third transduction was carried out at t=8h following a PBS wash and removal of 4-OHT. At t=24 h, cells were selected for shRNA-expression in 90 μg/mL hygromycin for 72 h. (**B**) Examples of the localization of the indicated Myc-tagged proteins (IF, cyan) and 53BP1 (IF, green) at telomeres detected with TTAGGG repeat FISH (red). DNA stained with DAPI. Merged and enlarged images are shown. Induction of Cre and AP-mediated FKBP dimerization as in (A). (**C**) Quantification of TIF response as detected in (B). Data represent means ±SDs of three biological replicates of 30 nuclei each. ns, P>0.05; *, P<0.05; **, P<0.01; ***, P<0.001; ****, P<0.0001. P values based on ordinary one-way ANOVA with multiple comparisons. (**D**) Detection of telomere fusions in metaphase spreads from the indicated cells. DNA stained with DAPI (red) and telomeres detected with FISH (green). (**E**) Quantification of telomere fusions represented in (D). Data represent means ±SDs of three biological replicates of 20 metaphase spreads each. ns, P>0.05; *, P<0.05; **, P<0.01; ***, P<0.001; ****, P<0.0001. P values based on ordinary one-way ANOVA with multiple comparisons. (**F**) Immunoblots for FKBP-TRF1 expressed in TRF2^F/-^ Lig4^-/-^ Cre-ER^T2^ MEFs detected with an antibody to TRF1. Cre was induced with 4-OHT, and TRF2 knockout was confirmed 96 h after 4-OHT addition with an antibody against mouse TRF2. +AP: addition of AP1903 at 72 h before harvest. +shTPP1: cells were infected thrice with a shTPP1 or shLuc retroviral vector. The second transduction was carried out at t=0 and coincided with 4-OHT addition. The third transduction was carried out at t=8h following a PBS wash and removal of 4-OHT. At t=24 h, cells were selected for shRNA-expression in 90 μg/mL hygromycin for 72 h. (**G**) Detection of telomere fusions in metaphase spreads from the indicated cells. DNA stained with DAPI (red) and telomeres detected with FISH (green). (**H**) Quantification of the percentage of telomeres that are fused detected as in (F). Data represent means ±SDs of three biological replicates of 30 nuclei each. ns, P>0.05; *, P<0.05; **, P<0.01; ***, P<0.001; ****, P<0.0001. P values based on ordinary one-way ANOVA with multiple comparisons. (**I**) Model of t-loop formation by a tetrameric TRF2 or TRF2 and depiction of how the lack of a free DNA end in the t-loop prevents activation of ATM signaling by MRN and blocks initiation of cNHEJ by Ku70/80.

## Discussion

Here we show that tetramerization of TRF2 is required for the protection of telomeres. Based on this finding, we generated a tetrameric version of TRF1 that can form t-loops in vivo and protect telomeres in cells devoid of TRF2. This orthogonal method for t-loop formation demonstrated that t-loops per se are sufficient for the repression of ATM signaling and cNHEJ.

It was proposed that t-loops would have the ability to repress cNHEJ, since the ring-shaped Ku70/80 heterodimer that initiates cNHEJ can only load on free DNA ends (Fig. 4I). However, the observation that reduced TRF2 expression induced ATM signaling at telomeres that remained resistant to cNHEJ and the finding that cNHEJ can be repressed by Rap1 had cast doubt on this view (*21–26*). Our interpretation is that TRF2 can employ two mechanisms to block cNHEJ. The primary block to cNHEJ is provided by t-loops making the telomere end inaccessible to Ku70/80, whereas Rap1-mediated inhibition of cNHEJ may function as a secondary mechanism to avoid joining of telomeres that are temporarily linear.

The data show that t-loops can repress ATM signaling even when TRF2 is absent. The ATM kinase is activated by MRN (Mre11, Rad50, Nbs1) when it recognizes a free DNA end (*49*). MRN undergoes a conformational change at a DNA end but how it activates the ATM kinase is unclear (*50*). Regardless of the mechanistic details, the constrained telomere end in the t-loop configuration will prevent MRN from activating ATM (Fig. 4I). Our data on the repression of ATM by t-loops in cells lacking TRF2 agrees with findings in mouse ES cells, where a TRF2-independent mechanism leads to t-loops and repression of ATM signaling (*27*, *28*). However, the factor(s) that generate t-loops in ES cells are not known and it is not excluded that the repression of ATM in this setting is t-loop independent. As is the case for the repression of cNHEJ, TRF2 possesses more than one mechanism repressing the formation of DNA damage foci at telomeres. First, it was shown that TIN2 contributes to the repression of ATM but the mechanism by which TIN2 acts is not known (*51*). Second, TRF2 can use its highly conserved iDDR module to reduce the DNA damage response at telomeres (*52*).

## Methods

### Cell culture and expression constructs

SV40LT-immortalized TRF2^F/F^ Rosa26-Cre-ER^T2^, TRF2^F/-^Lig4^-/-^ Rosa26- Cre-ER^T2^, and TRF2^F/F^ Ku70^-/-^ MEFs have been described previously (*10*, *11*, *53*). MEFs were cultured in Dulbecco’s Modified Eagle Medium (DMEM, Corning) supplemented with 10% fetal bovine serum (FBS) (Gibco), non-essential amino acids (Gibco), 2 mM l-glutamine (Gibco), 100 U/mL penicillin, and 100 μg/mL streptomycin (Gibco). Phoenix E cells (ATCC) used for retroviral packaging were cultured in DMEM supplemented with 10% bovine calf serum (BCS), non-essential amino acids, l-glutamine, and penicillin–streptomycin as above. Ku-deficient MEFs were infected with pMMP Hit & Run Cre thrice at 12-h intervals to introduce Cre recombinase, followed by media change as described previously (*11*). Cell harvest time points indicate hours after the second infection. Cells containing Rosa26-Cre-ER^T2^ were treated with tamoxifen (4-OHT, 1 μM) for 8 h to induce Cre recombinase expression, followed by a media change. Cell harvest time points indicate hours after tamoxifen addition.

All expression constructs were generated using Gibson assembly. Proteins were expressed constitutively using pLPC retroviral vector selected in 2-3 μg/mL puromycin, except for proteins in Figs. 2 and S3, which were expressed under the control of a dox-inducible promoter. MEFs that stably express rtTA after retroviral infection with pQCXIB-rtTA were selected in 5 μg/mL blasticidin. The MEFs were subsequently infected with pRetroX retroviral vector carrying dox-inducible constructs. Infected cells were selected in 2-3 μg/mL puromycin. Mouse TPP1 shRNA (GGACACATGGGCTGACGGA) (*54*) and luciferase shRNA (ACAACTTTACCGACCGCGCC) were introduced by 3 retroviral (pSUPER-Retro-Hygro) infections at 8-12 h intervals and selected in 90 μg/mL hygromycin. Drug treatments were as follows. FKBP dimerization: 100 nM AP1903 (ApexBIO B4168); ATR inhibition: 1 μM AZ20 (Selleck Chemical), 24 h; ATM inhibition: 5 μM KU55933 (Selleck Chemical), 24 h; PARP1 inhibition: 4 μM Olaparib (Selleck Chemicals), 120 h.

### Immunoblotting

Whole-cell extracts were prepared by resuspending cell pellets in RIPA lysis buffer (150 mM NaCl, 1% NP-40, 0.5% sodium deoxycholate, 0.1% SDS, 50 mM Tris-HCl, pH 8) containing protease inhibitors (100 μM PMSF + complete protease-inhibitor cocktail (Sigma P7626) and denatured in Laemmli buffer at 100^0^C for 5 min. Equal amounts of proteins as measured by Pierce™ BCA Protein Assay Kit (cat. #23225) were separated on 4-12% SDS-PAGE and transferred to nitrocellulose membranes. Blocking was performed with either 5% milk or LICOR Intercept (PBS) Blocking Buffer (LICOR cat. #927-70001). The following antibodies were used: mTRF2 (#1254, de Lange lab (*11*), 1:5000); mTRF1 (#1448, de Lange lab (*12*), 1:2000); Myc-tag (9B11, Cell Signaling Technology; 1:1,000); and γ-tubulin (GTU488, Sigma; 1:20,000). Primary antibodies were diluted in either 2% milk or LICOR Intercept (PBS) Blocking Buffer + 0.1% Tween and incubated, shaking, with the membrane at 4°C overnight. The following secondary antibodies were used: IRDye 800CW Goat anti-Rabbit IgG Secondary Antibody (LICOR cat. # 926-32211) and IRDye 800CW Goat anti-Mouse IgG Secondary Antibody (LICOR cat. # 926-32210). Secondary antibodies were incubated, shaking, with membranes for 30-60 min at room temperature. Imaging was performed using a GE Typhoon system.

### Immunofluorescence combined with FISH

IF-FISH was performed as described (*55*). Cells grown on coverslips were fixed for 10 min in 3% paraformaldehyde/2% sucrose at room temperature (RT) and permeabilized for 10 min in 0.5% Triton X-100 buffer (0.5% Triton X-100, 20mM Hepes-KOH, 50mM NaCl, 3mM MgCl_2_, 300mM sucrose). Coverslips were incubated in blocking buffer (1 mg/mL BSA, 2% goat serum, 0.1% Triton X-100, and 1mM EDTA pH 8 in PBS) for 30 min, then labeled with primary antibodies at RT for 1 h or overnight at 4^0^C. The antibodies used were 53BP1 (ab175933, Abcam) and Myc-tag (9B11, Cell Signaling). Coverslips were washed three times in 1X PBS and incubated for 30 min with secondary antibodies labeled with Alexa 488 or Alexa 555 (Molecular Probes; 1:1,000). Coverslips were washed with PBS, fixed again for 10 min in 3% paraformaldehyde/2% sucrose at RT, dehydrated in 70%, 90%, and 100% ethanol for 2 min each, and allowed to air dry. Hybridizing solution (70% formamide, 1 mg/mL blocking reagent (Roche), and 10 mM Tris-HCl, pH 7.2) containing Alexa 647-OO-(TTAGGG)_3_ PNA probe (PNA Bio, Newbury Park, CA) was added to each coverslip, the coverslips were heated for 10 min at 80^0^C on a heat block and incubated in the dark for 2 h at room temperature or overnight at 4^0^C. The coverslips were washed twice for 15 min with 70% formamide/10 mM Tris-HCl (pH 7.2) and three times for 5 min with PBS. The DNA was counterstained by including DAPI in the second PBS wash. Coverslips were mounted in ProLong Gold antifade (Sigma) and imaged on a DeltaVision (Applied Precision) equipped with a cooled charge-coupled device camera (DV Elite CMOS Camera), a PlanApo ×60, 1.42 NA objective (Olympus), and SoftWoRx software. Telomere dysfunction induced foci (TIF) were scored as described previously(*31*). Image analysis was performed using Fiji ImageJ (version 2.1.0/1.53e).

### Metaphase FISH

Metaphase FISH for telomeric DNA was performed as described previously with some modifications (*55*). Briefly, mitotic cells were arrested in metaphase with 2 μg/mL colcemid for 1 h before harvest. Cells were trypsinized, swollen in 0.075M KCl at 37^0^C for 10 min and fixed in 3:1 methanol:acetic acid. After dropping metaphase spreads, the slides were aged overnight. The following day, slides were rehydrated in 1X PBS, fixed in 4% paraformaldehyde, dehydrated in 70%, 90%, and 100% ethanol for 2 min each, and allowed to air dry. The slides were treated with a hybridizing solution (70% formamide, 1 mg/mL blocking reagent (Roche), and 10 mM Tris-HCl pH 7.2) containing a Cy3-OO-(TTAGGG)_3_ PNA probe (PNA Bio, Newbury Park, CA) and denatured at 80°C for 10 min on a heat block. The slides were washed twice with 70% formamide, 10 mM Tris-HCl pH 7.2 for 15 min each, and three times in PBS for 5 min each. The DNA was counterstained by including DAPI in the second PBS wash, dehydrated in 70%, 90%, and 100% ethanol for 2 min each, and allowed to air dry. Slides were mounted in ProLong Gold antifade (Sigma) and imaged on a DeltaVision (Applied Precision) equipped with a cooled charge-coupled device camera (DV Elite CMOS Camera), a PlanApo ×60, 1.42 NA objective (Olympus), and SoftWoRx software.

### t-loop assay

T-loop assay was performed as described previously (*17*) with minor modifications. Briefly, 10- 15 × 10^6^ cells were resuspended in fibroblast lysis buffer (12.5 mM Tris, pH 7.4, 5 mM KCl, 0.1 mM spermine, 0.25 mM spermidine, and 175 mM sucrose, supplemented with a protease-inhibitor cocktail (Roche)) at a concentration of 8 × 10^6^ cells/mL and equilibrated on ice for 10 min. Cells were lysed by adding NP-40 to a final concentration of 0.2% and incubating on ice for 5 min. Nuclei were spun down at 1,000 <u>g</u> for 5 min, washed in 10 mL ice-cold nuclei wash buffer (10 mM Tris-HCl pH 7.4, 15 mM NaCl, 60 mM KCl, 5 mM EDTA, and 300 mM sucrose), and resuspended in 3 mL nuclei wash buffer. The nuclei were treated with 10 µg/mL of trioxsalen while on ice for 5 min in the dark with constant stirring. Cross-linking was initiated under 365 nm UV light, while the nuclei were still on ice with constant stirring for 5 min in the dark. The process of incubation and cross-linking was repeated once. The isolated nuclei were washed with nuclei wash buffer and then suspended in a solution of 50% glycerol and 50% nuclei wash buffer at a concentration of 2 × 10^7^ nuclei/mL. Subsequently, nuclei were diluted 1:10 in spreading buffer (10 mM Tris-HCl, pH 7.4, 10 mM EDTA, 0.1% SDS, and 1 M NaCl) and lysed by gentle pipetting. DNA was cytospun onto coverslips (Zeiss, cat. No. 474030-9000-000). The standard FISH procedure (see above) was used to hybridize an Alexa 488-OO-(TTAGGG)_3_ PNA probe and DNA spreads were then imaged on a GE OMX V4 microscope. At least 100 countable molecules were scored per sample. These molecules were divided into two distinct categories - “linear” and “loops”. As detailed previously (*17*), linear molecules were uniform, continuous, largely straight molecules with no branches, while loops were continuous molecules with a visible hollow loop with an aperture of at least 0.01 μm^2^ at one end. Any molecules that could not be classified as either linear or loops were excluded from the analysis.

### Purification of E. coli expressed proteins

N-terminally His6-Smt3-tagged proteins (pRSFDuet-1, Novagen) were expressed in Rosetta (DE3) competent cells (Novagen) grown in LB Broth (Miller, Research Products International) at 37°C to an OD600 of 0.4. The temperature was lowered to 18°C while the cells continued growing to an OD600 of 0.6, and expression was induced with 0.5 mM IPTG for 16 h at 18°C. Cells were collected by centrifugation at 2,500 g, resuspended in a buffer containing 20 mM Tris-HCL pH 8, 500 mM NaCl, 4 mM β-mercaptoethanol (β-ME), 20 mM imidazole, and 10% glycerol, supplemented with complete EDTA-free protease inhibitor cocktail (Roche), and flash frozen in liquid nitrogen. Cells were thawed in cold water and lysed with 2% v/v Triton X-100, 50 mM Tris-HCl pH 8, 500 mM NaCl, 15 mM β-ME, 20 mM imidazole-HCl pH 7.5, 40 μg/mL lysozyme (Sigma), 10 μg/mL DNaseI (Sigma), 1 mM CaCl2, and 1 mM phenylmethylsulfonyl fluoride (PMSF). Lysates were sonicated and clarified by centrifugation at 40,000 g at 4°C for 45 min. The supernatant was incubated for 1 h with Ni-NTA resin (Invitrogen) equilibrated in buffer containing 20 mM Tris-HCL pH 8, 500 mM NaCl, 4 mM β-ME, and 20 mM imidazole and washed with 20-50 CV of the same buffer. Bound protein was eluted in a buffer containing 20 mM HEPES-NaOH pH 7.5, 150 mM NaCl, 0.5 mM tris(2-carboxyethyl)phosphine (TCEP), 250 mM imidazole, and 5% glycerol. The His6-Smt3 tag was digested with His6-Ulp1 protease for 3 h at 8°C, and the products were incubated with Ni-NTA to remove His6-Smt3, His6-Ulp1, and uncleaved protein. The flowthrough was concentrated to 500 μL using a 10- or 30-kDa cutoff Amicon concentrator (Millipore) and loaded onto a Superdex 200, Superdex 75, or Superose 6 Increase 10/300 GL column (Cytiva) equilibrated in 20 mM HEPES-NaOH pH 7.5, 300 mM NaCl, 0.5 mM TCEP-NaOH. Due to insolubility, His6-Smt3-TRF2ΔB-F59A and His6-Smt3- TRF2ΔBΔHinge-F59A were prepared from inclusion bodies and renatured. Cell pellets were thawed in cold water and added to 20 mL of cold 20 mM Tris-HCl pH 8.0 and 500 mM NaCl, sonicated, and clarified by centrifugation at 10,000 g at 4°C for 15 min. Pellets were resuspended in 10 mL ice-cold 50 mM Tris-HCl pH 8.0, 0.2 mM PMSF, and 1% Triton X-100. The lysate was clarified as above, and the pellet was resuspended in 5 mL of ice-cold 50 mM Tris-HCl pH 8.0 and 0.2 mM PMSF. After repeating this step once more, the final pellet was resuspended in 5 mL of ice-cold 50 mM Tris-HCl pH 8, 8 M urea, 10 mM dithiothreitol (DTT), and 0.2 mM PMSF, and the suspension was rotated at 8°C for 2 h. The denatured protein solution was clarified by centrifugation at 30,000 g at 4°C for 30 min. 15 mg of the denatured protein mixture was added to 62.5 mL of 3 M urea, 100 mM Tris-HCl pH 8, 0.4 M L-arginine monohydrochloride, and 20 mM reduced L-glutathione. This mixture was incubated for 24 h at 8°C with stirring at 500 rpm. The protein mixture was renatured by dialysis in 15 kDa MW cutoff dialysis tubing (Spectro Laboratories, cat. No. 25223-923) in 2 L 20 mM Tris-HCl pH 8, 500 mM NaCl at 8°C for 8 h with continuous stirring and buffer changes every 2 h, followed by an additional 16 h without stirring. The His6-Smt3 tags were digested with His6-Ulp1 protease for 3 h at 8°C and removed using Ni-NTA resin as described above. The flowthrough was concentrated using an Amicon concentrator as described above. Protein concentration was measured on a Nanodrop1000 spectrophotometer. All proteins were flash frozen in liquid nitrogen and stored in aliquots at -80°C.

### In vitro transcription and translation

Myc-tagged genes were cloned into pTNT vector (Promega L5620). Proteins were expressed using TnT® Quick Coupled Transcription/Translation System (Promega L1170), according to manufacturer’s instructions and stored at -80°C.

### Electrophoresis Mobility Shift Assay

For EMSAs, the double-stranded DNA probe with 4 telomeric repeats were synthesized as two single-stranded oligonucleotides (Invitrogen). Oligo 1 - 5’- GTACCCGGGGATCGTCGACTCTAGAGGGGCCCTAACCCTAACCCTAACCCTAACCCGGGCTCGAATTCGATCCTCTAGAGTCGACCTGCAGGCATGCA-3’; Oligo 2 - 5’- AGCTTGCATGCCTGCAGGTCGACTCTAGAGGATCGAATTCGAGCCCGGGTTAGGGTTAGGGTTAGGGTTAGGGCCCCTCTAGAGTCGACGATCCCCGG-3’. The 2 oligos were annealed and fluorescently labeled with Cy5.5-dUTP using Klenow DNA polymerase. Klenow fill-in was performed at 37^0^C for 45 min in a 25 μL reaction with 150 ng of annealed probe DNA (628 μM), 60 μM of dATP, dCTP, dGTP, and Cy5.5 labeled dUTP (Lumiprobe), 1X NEB Buffer 2, and 5 U/mL Klenow exo^-^ (NEB). Klenow was heat-inactivated and 5 μg E. coli DNA (Sigma D-2001, type VIII), sheared as described (*56*), was added to reduce non-specific binding by TRF2. The buffer was exchanged twice using Biogel P-30 spin columns to remove free dNTPs. The protein-DNA binding reaction was carried out at room temperature for 20 min. The reaction mixture (4% glycerol, 50 ng/mL sheared E. coli DNA, 150 mM Tris-HCl pH 7.5, 100 ng/mL β-casein, 0.5 nM probe, 1 mM EDTA, and 100 mM NaCl) was incubated with the indicated concentrations of purified TRF2 proteins or 2 μL of RRL mixture. The protein-DNA complexes were resolved on 0.8% agarose gels in 0.1x TBE run in 0.1x TBE buffer at 200 V for 40 min at room temperature. Imaging was performed using a GE Typhoon system. Image analysis and quantification was performed using Fiji ImageJ (version 2.1.0/1.53e).

### Quantification and Statistical Analysis

The number of biological replicates performed is indicated in each figure and corresponds to the number of data points plotted. All metaphase FISH and IF-FISH images were scored manually using FIJI ImageJ (version 2.1.0/1.53e). Statistical analysis was performed in GraphPad Prism 9.4.1.

## Materials

### Cell Lines

| Line | Species | Source/reference |
| --- | --- | --- |
| Phoenix E | *Homo sapiens* | ATCC CRL-3213 |
| HEK293FT | *Homo sapiens* | ThermoFisher R70007 |
| TRF2<sup>F/F</sup> Cre-ER<sup>T2</sup> MEFs | *Mus musculus* | (11) |
| TRF2<sup>F/-</sup> Lig4<sup>-/-</sup> Cre-ER<sup>T2</sup> MEFs | *Mus musculus* | (11) |
| TRF2<sup>F/F</sup> Ku70<sup>-/-</sup> MEFs | *Mus musculus* | (10) |

### Plasmids

| Plasmid | Source/reference | Gene Species |
| --- | --- | --- |
| pLPC vector | Addgene 12540 | n/a; vector |
| pLenti-sgRNA | Addgene 71409 | n/a; vector |
| pRetroX vector | Takara Bio 631188 | n/a; vector |
| pRSF-Duet vector | Millipore Sigma<br>71341-3 | n/a; vector |
| pTNT vector | Promega L5620 | n/a; vector |
| pLPC Myc-mTRF1 | Cloned here | *Mus musculus* |
| pLPC Myc-mTRF2 | Cloned here | *Mus musculus* |
| pLPC Myc-mTRF2-Topless | Cloned here | *Mus musculus* |
| pLPC Myc-FKBP(V36)-mTRF2-Topless | Cloned here | *Mus musculus* |
| pLPC Myc-FKBP(V36)-mTRF1 | Cloned here | *Mus musculus* |
| pLPC Myc-FKBP(V36)-mTRF1ΔTRFH | Cloned here | *Mus musculus* |
| pRetroX Myc-mTRF2 | Cloned here | *Mus musculus* |
| pRetroX Myc-mTRF2ΔHinge | Cloned here | *Mus musculus* |
| pRetroX Myc-mFKBP(V36)-TRF2ΔHinge | Cloned here | *Mus musculus* |
| pLPC Myc-FKBP(WT)-TRF1 | Cloned here | *Mus musculus* |
| pRSF-Duet His(6x)-Sumo-TRF2ΔBasic | Cloned here | *Mus musculus* |
| pRSF-Duet His(6x)-Sumo-TRF2ΔBasic-F59A | Cloned here | *Mus musculus* |
| pRSF-Duet His(6x)-Sumo-TRF2ΔBasicΔTRFH | Cloned here | *Mus musculus* |
| pRSF-Duet His(6x)-Sumo-TRF2ΔBasicΔHinge | Cloned here | *Mus musculus* |
| pRSF-Duet His(6x)-Sumo-TRF2ΔBasicΔHinge-F59A | Cloned here | *Mus musculus* |
| pRSF-Duet His(6x)-Sumo-MBP-mTRF2 Myb | Cloned here | *Mus musculus* |
| pTNT-Duet Myc-TRF2ΔBasic | Cloned here | *Mus musculus* |
| pTNT-Duet Myc-TRF2ΔBasic F59A | Cloned here | *Mus musculus* |
| pTNT-Duet Myc-TRF2ΔBasicΔTRFH | Cloned here | *Mus musculus* |
| pTNT-Duet Myc-TRF2ΔBasicΔHinge | Cloned here | *Mus musculus* |
| pTNT-Duet Myc-TRF2ΔBasicΔHinge-F59A | Cloned here | *Mus musculus* |
| pTNT-Duet Myc-TIN2 TRFH-TRF2 Myb | Cloned here | *Mus musculus* |

### Plasmid sequencing primers

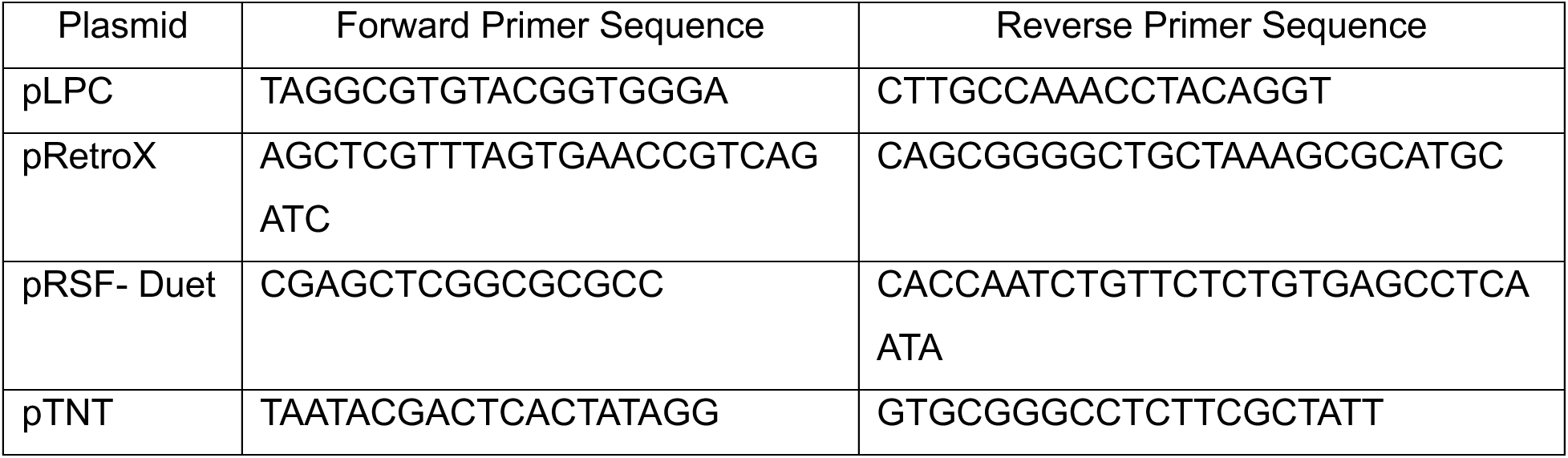

### Antibodies

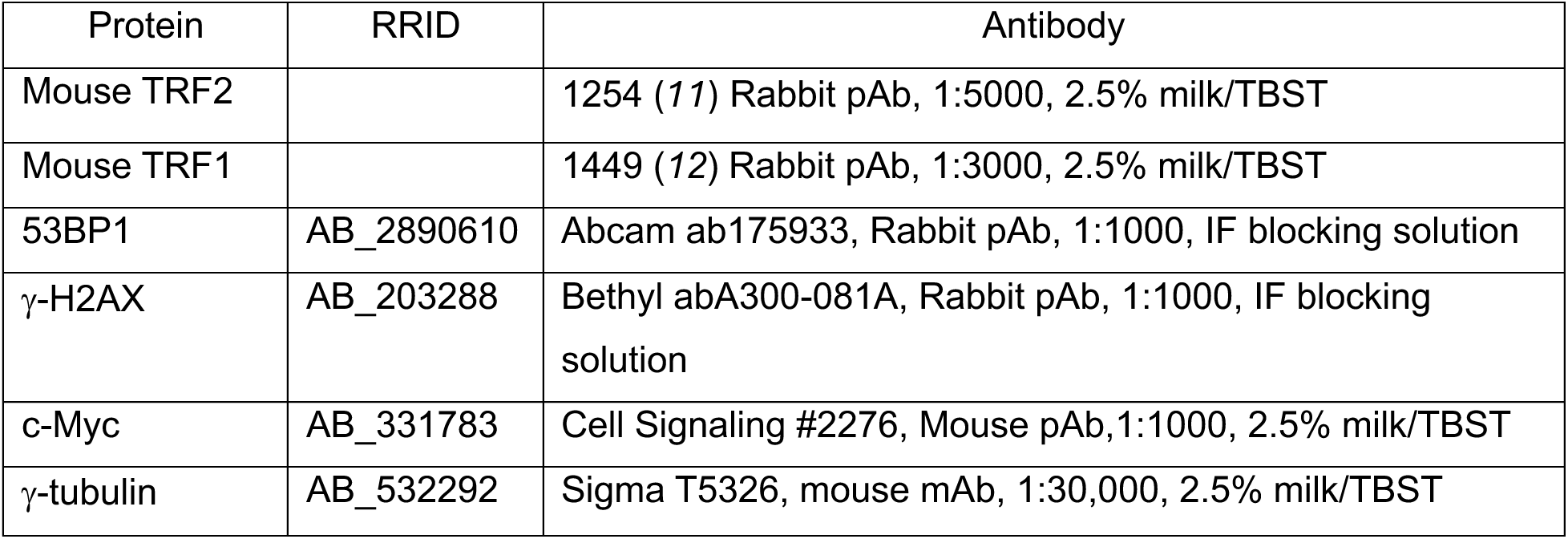

## Acknowledgements

We thank H. Moss for piloting the EMSA experiments and identifying the unusual binding pattern of the F59A mutant which led us to consider tetramerization. We thank members of the de Lange laboratory for helpful discussion of this work. We thank members of the Rockefeller University Bio-Imaging Resource Center, especially A. North, for assistance with t-loop imaging. This study was supported by grants from the National Cancer Institute R35CA210036 (T.d.L.), the National Institute on Aging PHSR01AG016642 (T.d.L.) and the National Science Foundation GRFP2020291008 (A.G.).

## Supplementary

**Fig. S1.**
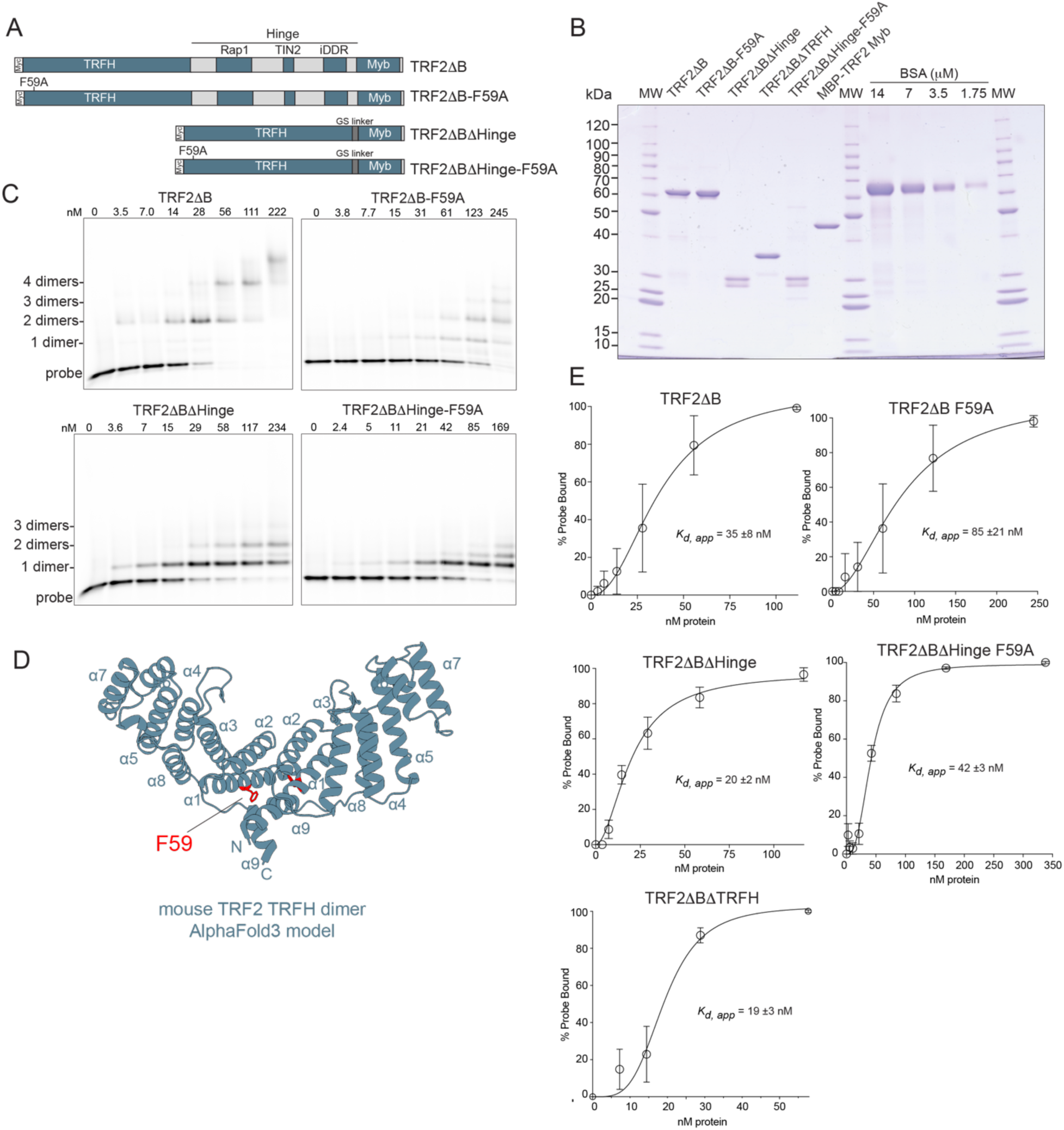
TRF2 binds telomeric DNA as a tetramer. (**A**) Schematics of TRF2τιB (lacking the Basic domain), TRF2τιB-F59A, TRF2τιBτιHinge, and TRF2τιBτιHinge-F59A. **(B)** Coomassie gel of indicated TRF2 proteins schematized in (A) purified from E. coli. BSA standard was used to verify protein concentrations. 2μl was loaded per lane. The doublet observed in TRF2ΔBΔHinge and TRF2ΔBΔHinge-F59A is consistently observed in protein purified from E.coli, synthesized by in vitro coupled transcription/translation, and in protein extracts from MEFs expressing TRF2ΔBΔHinge in vivo. The reason for the doublet is not known. **(C)** EMSAs with the indicated TRF2 proteins schematized in (A) with indicated protein concentrations. The probe concentration was 0.5 nM. Note that the smallest complex formed by TRF2ΔB is consistent with a tetramer (two dimers) whereas the smallest complex formed by TRF2ΔB-F59A represents a dimer. **(D)** Structure of TRF2’s TRFH dimerization interface with F59 residue indicated in red, modeled by AlphaFold3. **(E)** Affinity curves for EMSAs in (C), calculated % probe bound from three technical replicates. Specific binding with Hill slope, error bars reflect mean and standard deviation.

**Fig. S2.**
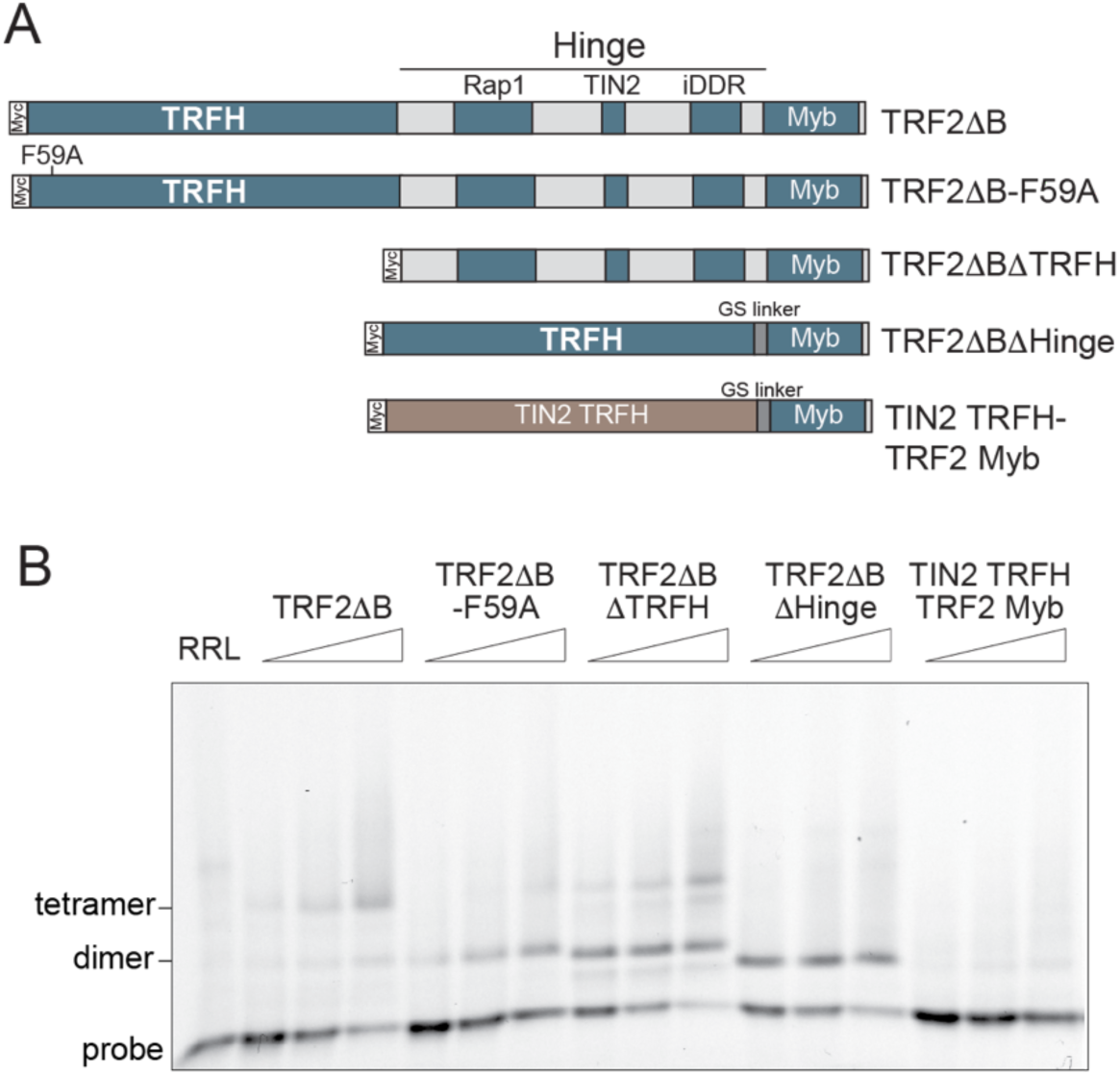
Confirmation of tetrameric DNA binding by TRF2. (**A**) Schematics of Myc-tagged TRF21′B, TRF21′B-F59A, TRF21′B1′TRFH, TRF21′B1′Hinge, and TIN2 TRFH-TRF2 Myb. The TRFH of TIN2 does not dimerize. **(B)** EMSA with the indicated TRF2 proteins schematized in (A) produced by coupled in vitro transcription/translation in RRL. Lanes represent gel-shifts with 0.5, 1.0, and 2.0 μl of RRL. The monomeric TIN2 TRFH-TRF2 Myb fusion protein does not show detectable DNA binding activity. The other binding patterns are consistent tetrameric binding by TRF2.

**Fig. S3.**
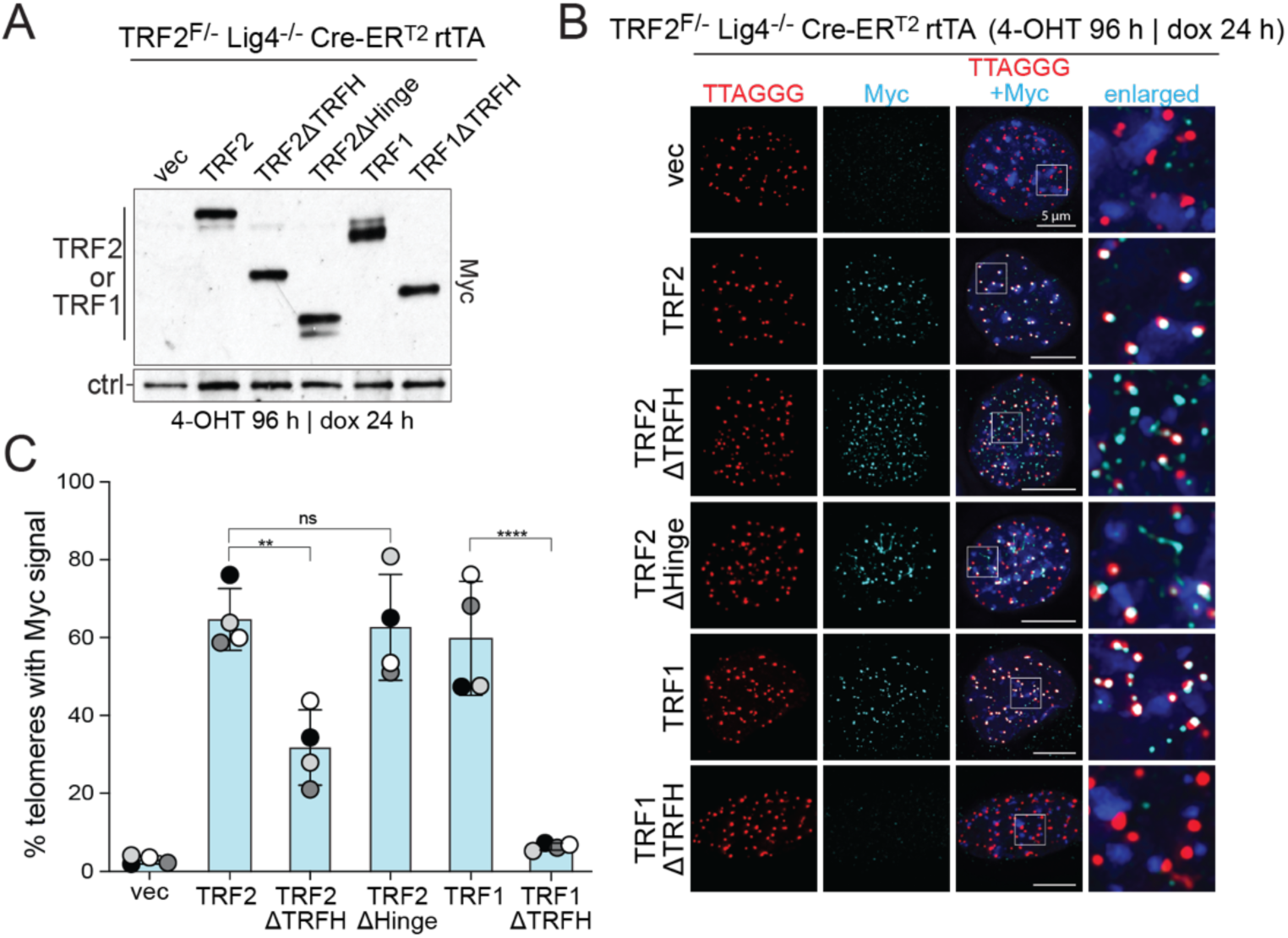
Telomeric localization of TRF21′TRFH but not TRF11′TRFH. (**A**) Immunoblots for the indicated Myc-tagged TRF2 and TRF1 proteins expressed in TRF2^F/-^ Lig4^-/-^ Cre-ER^T2^ MEFs detected with an antibody to Myc. Relatively equal levels of proteins were expressed from a Dox-inducible promoter by treating with different concentrations of doxycycline (TRF2 and TRF21′TRFH: no Dox; TRF21′Hinge: 2 μg/mL; TRF1: 10 ng/mL, TRF11′TRFH: 10 ng/mL). Cre was induced with 4-OHT for 96 h. Doxycyclin was added 24 h before harvest. **(B)** Examples of the localization of the indicated Myc-tagged proteins (IF, cyan) at telomeres detected with TTAGGG repeat FISH (red). DNA stained with DAPI. Merged and enlarged images are shown. **(C)** Quantification of the percentage of telomeres with Myc signal colocalization as detected in **(D)** Data represent means ±SDs of three biological replicates with 20 metaphases each. ns, P>0.05; *, P<0.05; **, P<0.01; ***, P<0.001; ****, P<0.0001. P values based on ordinary one-way ANOVA with multiple comparisons.

**Fig. S4.**
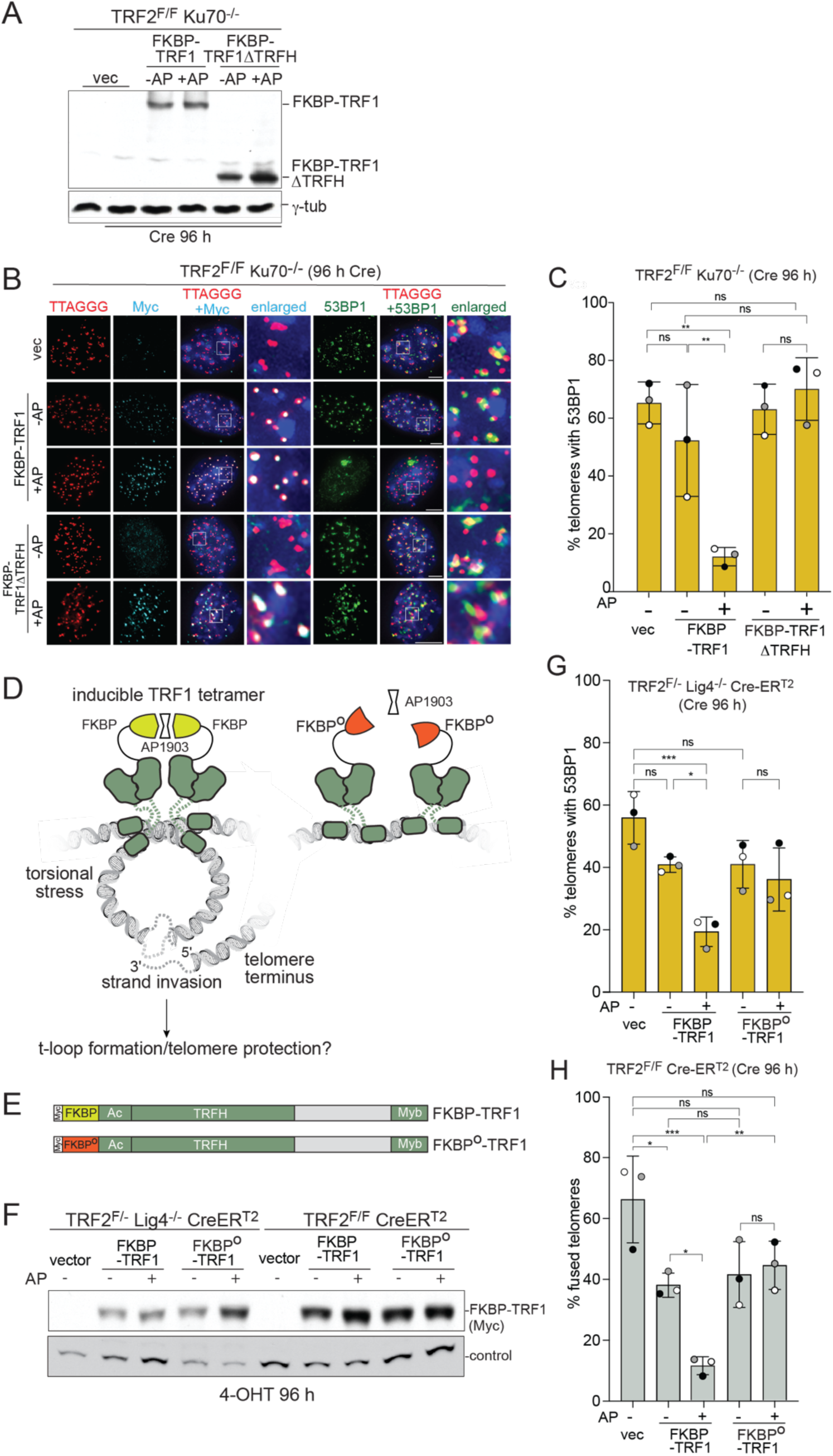
Confirmation of telomeric protection by tetramerized FKBP-TRF1. (**A**) Immunoblots for FKBP-TRF1 and FKBP-TRF11′TRFH proteins expressed in TRF2^F/-^ Ku70^-/-^ MEFs detected with an antibody to Myc. Cre was induced with 4-OHT for 96 h. +AP: addition of AP1903 at 72 h prior to harvest. To achieve equal induction, the following dox concentrations were used: TRF2, 0 nM; TRF21′TRFH, **(B)** Examples of the localization of the indicated Myc-tagged proteins (IF, cyan) and 53BP1 (IF, green) at telomeres detected with TTAGGG repeat FISH (red). DNA stained with DAPI. Merged and enlarged images are shown. Induction of Cre and AP-mediated FKBP dimerization as in (A). **(C)** Quantification of the percentage of telomeres with 53BP1 signal colocalization as detected in (B). Data represent means ±SDs of three biological replicates of 30 nuclei each. ns, P>0.05; *, P<0.05; **, P<0.01; ***, P<0.001; ****, P<0.0001. P values based on ordinary one-way ANOVA with multiple comparisons. **(D)** Model of t-loop formation by tetrameric TRF1 as dependent on induced dimerization of FKBP by AP1903. FKBP^0^ does not bind to AP1903 and therefore fails to dimerize in the presence of the drug. **(E)** Schematics of Myc-tagged FKBP-TRF1 and FKBP^0^-TRF1 expressed in MEFs. **(F)** Immunoblots for FKBP-TRF1 and FKBP^0^-TRF1 proteins expressed in TRF2^F/-^ Lig4^-/-^ Cre-ER^T2^ and in TRF2^F/-^ Cre-ER^T2^ MEFs detected with an antibody to Myc. Cre was induced with 4-OHT for 96 h. +AP: addition of AP1903 at 72 h prior to harvest. Asterisk: nonspecific band. **(G)** Quantification of the percentage of telomeres colocalizing with 53BP1. Data represent means ±SDs of three biological replicates of 30 nuclei each. ns, P>0.05; *, P<0.05; **, P<0.01; ***, P<0.001; ****, P<0.0001. P values based on ordinary one-way ANOVA with multiple comparisons. **(H)** Quantification of the percentage of telomeres that are fused. Data represent means ±SDs of three biological replicates of 20 metaphase spreads each. ns, P>0.05; *, P<0.05; **, P<0.01; ***, P<0.001; ****, P<0.0001. P values based on ordinary one-way ANOVA with multiple comparisons.

**Fig. S5.**
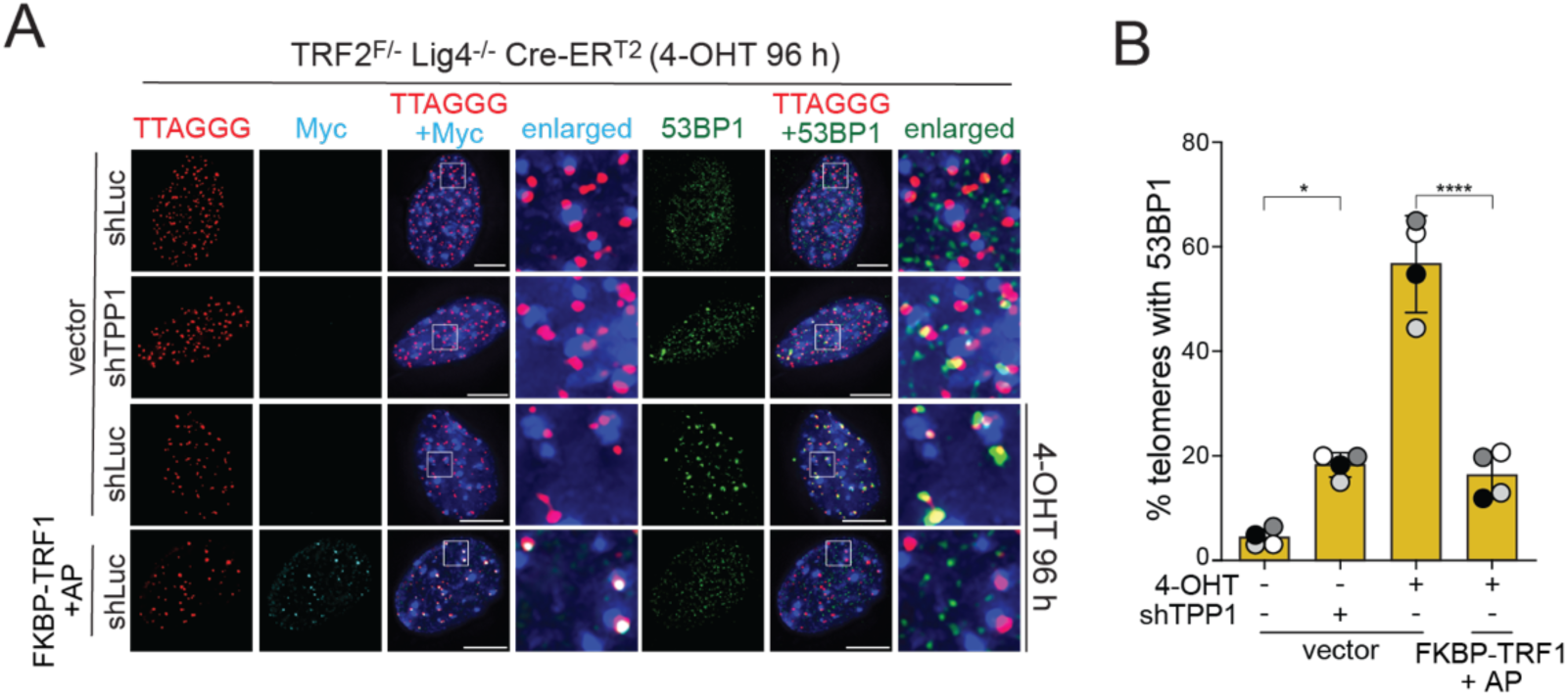
TIF assay to verify that shTPP induces a telomere damage signal in Lig4^-/-^ cells. (**A**) Examples of the localization of the indicated Myc-tagged proteins (IF, cyan) and 53BP1 (IF, green) at telomeres detected with TTAGGG repeat FISH (red) in TRF2^F/-^ Lig4^-/-^ Cre-ER^T2^ MEFs. DNA stained with DAPI. Merged and enlarged images are shown. Cre was induced with 4-OHT for 96 h. +AP: addition of AP1903 at 72 h prior to harvest. +shTPP1: cells were infected thrice with a retrovirus expressing shTPP1 or shLuc. The second transduction was carried out at t=0 and coincided with 4-OHT addition. The third transduction was carried out at t=8h following a PBS wash and removal of 4-OHT. At t=24 h, cells were selected for shRNA-expression in 90 μg/mL hygromycin for 72 h. **(B)** Quantification of the percentage of telomeres colocalizing with 53BP1 signals. Data represent means ±SDs of three biological replicates of 30 nuclei each. ns, P>0.05; *, P<0.05; **, P<0.01; ***, P<0.001; ****, P<0.0001. P values based on ordinary one-way ANOVA with multiple comparisons.

